# Canonical Wnt Signaling Induces Focal Adhesion and Integrin Beta-1 Endocytosis

**DOI:** 10.1101/2022.01.11.475934

**Authors:** Nydia Tejeda-Muñoz, Marco Morselli, Yuki Moriyama, Pooja Sheladiya, Matteo Pellegrini, Edward M. De Robertis

## Abstract

During canonical Wnt signaling, the Lrp6 and Frizzled co-receptors bind to the Wnt growth factor and the complex is endocytosed and sequestered together with Glycogen Synthase Kinase 3 (GSK3), Dishevelled (Dvl), and Axin inside the intraluminal vesicles of late endosomes, known as multivesicular bodies (MVBs). Here we present experiments showing that Wnt causes the endocytosis of focal adhesion (FA) proteins and depletion of Integrin β 1 (ITGβ1) from the cell surface. FAs and integrins link the cytoskeleton to the extracellular matrix. Wnt-induced endocytosis caused ITGβ1 depletion from the plasma membrane and was accompanied by striking changes in the actin cytoskeleton. In situ protease protection assays in cultured cells showed that ITGβ1 was sequestered within membrane-bounded organelles that corresponded to Wnt-induced MVBs containing GSK3 and FA-associated proteins. An *in vivo* model using *Xenopus* embryos dorsalized by *Wnt8* mRNA showed that ITGβ1 depletion decreased Wnt signaling. The finding of a crosstalk between two mayor signaling pathways, canonical Wnt and focal adhesions, should be relevant to human cancer and cell biology.

## Introduction

Recent work has shown a fundamental role for membrane trafficking in the cell biology of canonical Wnt signaling (Albrecht et al., 2021) in addition to the well-established transcriptional effects mediated by β-catenin stabilization (Nusse and Clevers, 2017). Binding of the Wnt growth factors to the Frizzled (Fz) and LDL-receptor related protein 6 (LRP6) co-receptors causes the formation of signalosomes at the plasma membrane and local inhibition of GSK3 (Bilić et al., 2007; Niehrs, 2012). The receptor complex is then endocytosed together with other cytoplasmic proteins such as Dishevelled (Dvl), Axin1, and GSK3 (Taelman et al., 2010; Kim et al., 2015). The local decrease of GSK3 activity in signalosomes rapidly triggers macropinocytosis and endocytosis of the receptor complex, GSK3, and Axin1 (Tejeda-Muñoz et al., 2019). These are rapidly trafficked into late endosomes and, via the Endosomal Sorting Complexes Required for Transport (ESCRT) machinery, into the intraluminal vesicles of multivesicular bodies (MVBs) (Taelman et al., 2010; Dobrowolski et al., 2012; Albrecht et al., 2018). The sequestration of GSK3 and Axin is necessary for the stabilization of β-catenin (Taelman et al., 2010). The decrease of GSK3 in the cytosol results in the stabilization of many other GSK3 substrate proteins in a novel pathway known as Wnt Stabilization Of Proteins (Wnt-STOP) (Kim et al., 2009; Taelman et al., 2010; Acebron et al., 2014).

Within a few minutes of Wnt addition of Wnt3a, endocytic vesicles are formed (Tejeda-Muñoz et al., 2019; Albrecht et al., 2020). These vesicles correspond to MVBs and contain GSK3 that becomes protected from proteinase K digestion inside membrane-bounded organelles after permeabilizing the plasma membrane with digitonin (Albrecht et al., 2018). Wnt treatment triggers a large increase of non-receptor mediated endocytosis of molecules such as bovine serum albumin (BSA) and high molecular weight Dextran (Albrecht et al., 2018; Tejeda et al., 2019), consistent with the view that Wnt signaling induces sustained macropinocytosis. Macropinocytosis is a non-receptor-mediated actin-driven process resulting from activation of p21-activated kinase-1 (Pak1) (Doherty and McMahon, 2009). This cell drinking mechanism leads to the formation of plasma membrane ruffles, actin tent poles, and macropinocytic cups that internalize extracellular fluid in vesicles of sizes 200 nm to 5 µm (Condon et al., 2018; Swanson and King, 2019). Macropinocytosis is driven by mechanisms very similar to those of phagocytosis, an ancient process that frequently involves integrins (Mylvaganam et al., 2021). On the other hand, receptor-mediated endocytosis leads to the formation of small vesicles of less than 100 nm visible only by electron microscopy mediated by the clathrin or caveolin machinery in a process designated micropinocytosis (Nichols and Lippincott-Schwartz, 2001; Doherty and McMahon, 2009). In the case of Wnt-induced macropinocytosis, internalized macromolecules are rapidly trafficked into lysosomes where they are degraded, leading to changes in metabolism (Albrecht et al., 2020).

The starting point of the present investigation came when we noticed that the large cytoplasmic vesicles induced by Wnt3a were formed in the periphery of cultured cells, mainly at the leading edge. This is also the location of focal adhesion (FA) proteins which mediate the interaction between the cytoskeleton and the extracellular matrix (ECM) via integrins (Abercrombie et al.,1971; Bachir et al., 2017). This localization reminded us of intriguing results presented by Capelluto et al., (2002) in which they described that overexpressed Dvl protein, which is also a component of Wnt MVBs, was found in actin cables and cytoplasmic puncta. Although not specified at the time, in their images the puncta were frequently at the tip of actin cables which is also the location of FAs (Capelluto et al., 2002). This led us to hypothesize that FAs and integrins might crosstalk with Wnt signaling.

In the present paper we show that within 20 minutes of Wnt3a addition, human fibroblasts and cultured cell lines underwent major rearrangements of the actin cytoskeleton and focal adhesions. Using cell surface biotinylation we showed that Wnt3a treatment leads to the internalization of Integrin β1 (ITGβ1). Proteinase K protection studies in situ revealed that ITGβ1 and other FA proteins, such as Zyxin and Src, were sequestered in MVBs after Wnt3a treatment. Finally, a sensitized *Xenopus* embryo assay showed that ITGβ1 depletion decreased phenotypes caused by overexpressed *Wnt8* mRNA, and that this effect was rescued by human *ITGβ1* mRNA. Taken together, the results suggest an unexpected connection between focal adhesions, integrins, and canonical Wnt signaling.

## Results

### Wnt3a causes a major rearrangement of the actin cytoskeleton and focal adhesions in human corneal stromal (HCSF) fibroblasts

Wnt signaling causes the uptake of large amounts of extracellular fluid by macropinosomes (Redelman-Sidi et al., 2018; Tejeda-Muñoz et al., 2019). In addition, a gene ontology analysis of biotinylated Wnt-induced Lrp6-APEX2 target proteins reported that interacting proteins were involved in actin cytoskeleton remodeling (Colozza et al., 2020). To examine whether Wnt induced changes in the cytoskeleton, primary human corneal stromal (HCSF) fibroblasts (Gallego-Muñoz et al., 2018) were treated with Wnt3a. Fibroblasts showed abundant vinculin FA sites at the tip of F-actin cables, particularly in the leading edge of the cell (Figure 1A-A’’). Wnt treatment caused the loss of adhesion sites within 20 min (Figure 1B-B’’). This indicated that the cytoskeleton responds to Wnt stimulation, which causes the loss of FAs.

**Figure 1.**
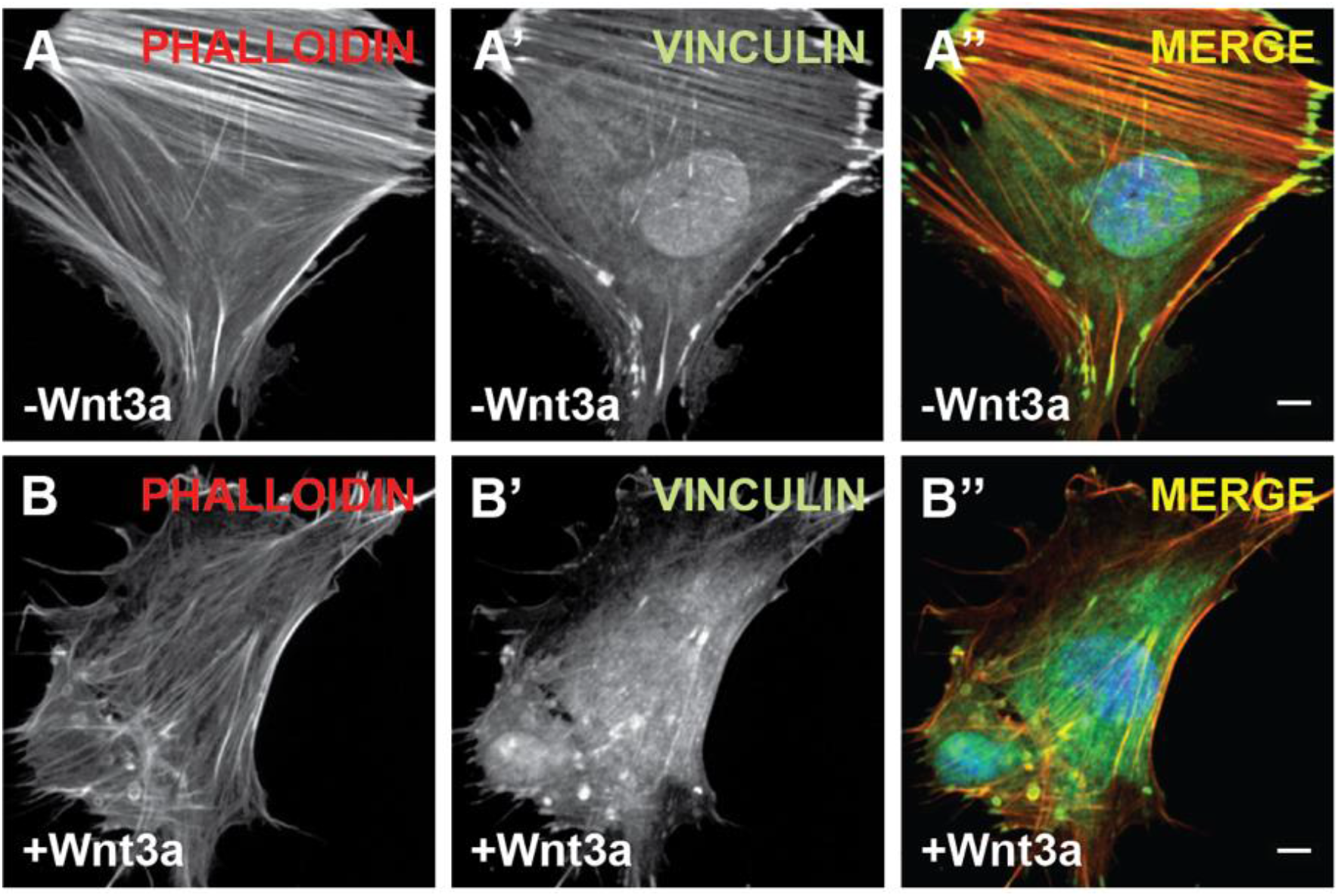
Wnt3a treatment for 20 minutes causes a major rearrangement of the actin cytoskeleton and focal adhesions in Human corneal stromal (HCSF) fibroblasts. (*A-A’’*) Image of HCSF fibroblast showing that the cells attach well to fibronectin-coated plastic, have prominent F-actin cables stained by Phalloidin (in red) and abundant focal adhesion sites immunostained with vinculin (in green). (*B-B’’*) Wnt3a treatment causes disorganization of in the cytoskeleton, vinculin becomes associated with intracellular vesicles sometimes surrounded by F-actin, and few focal adhesions are visible. Similar results were obtained in three independent experiments. Nuclei were stained with DAPI. Scale bars, 10 μm. **Figure supplement 1.** Macropinocytosis in different cellular models. **Figure supplement Video 1.** Mutation in Axin1 causes extensive membrane ruffling and macropinocytosis in HCC cells. **Figure supplement Video 2.** Mimicking Wnt with the GSK3 inhibitor LiCl triggers macropinocytic cup formation in cultured *Xenopus* animal cap cells. **Figure supplement Video 3.** Activated Ras-GFP triggers macropinocytosis.

Cancer cells undergo certain fundamental changes in terms of cell physiology to attain a malignant phenotype. Loss of cell-cell adhesion is caused by changes in the expression of adhesion proteins, which plays an important role in infiltration and metastasis (Bachir et al., 2017). To investigate how the Wnt pathway affects cell adhesion and motility, we used Hepatocellular carcinoma (HCC) Alexander1±Axin1 cells (Albrecht et al., 2020). This HCC cell line has a GSK3-binding region deletion in the tumor suppressor Axin1 that results in constitutive activation of the Wnt pathway. A stable cell line derivative that restores full-length Axin1 at physiological levels (Albrecht et al., 2020) serves as a control for the effect of Axin1 mutation. We filmed cells by differential interference contrast (DIC) light microscopy and found that mutation of Axin1 dramatically affected membrane motility, consistent with an increase in macropinocytosis (*Figure 1-figure supplement 1A*, *Video supplement 1*). This was manifested as wave-like extensions of the lamellipodium and extensive ruffling of the plasma membrane (*Video supplement 1*). When Axin1 was restored, membrane ruffling was lost (*Figure 1-figure supplement 1A’, Video supplement 1*). In a similar way, dissociated *Xenopus* animal cap cells injected with membrane GFP and Lifeact mRNA (marking F-actin) and cultured on fibronectin showed macropinocytic membrane ruffles within 20 min of addition of LiCl, a treatment that mimics Wnt signaling by inhibiting GSK3 (*Figure 1-figure supplement 1B-B’, Video supplement 2*). These circular ruffle membrane movements were very similar to those caused by transfection of oncogenic HRas (G12V mutation), a classical inducer of macropinocytosis (C. Ramirez et al. 2019) in transfected 3T3 fibroblasts (*Figure 1-figure supplement 1C-C’, Video supplement 3*).

The results indicate that Wnt signaling has major effects on the actin cytoskeleton, leading to loss of focal adhesions. This is accompanied by extensive membrane ruffling activity characteristic of macropinocytosis.

### Canonical Wnt signaling induces membrane vesicles at focal adhesions

Capelluto et al. reported that the DIX domain of Dishevelled is associated with actin cables and vesicular structures (Capelluto et al., 2002). We found it interesting that the membrane vesicles formed preferentially at the ends of actin cables, and hypothesized these might correspond to FAs. When HeLa cells were treated with Wnt3a for 20 min, puncta visible by DIC were formed at the leading edge of the cell (Figure 2A-D’, indicated with arrowheads). These vesicles concentrated endogenous GSK3, a protein normally uniformly distributed in the cytosol (Figure 2A’ and B’). Importantly, the Wnt-induced vesicles contained the FA protein Zyxin (Beckerle, 1986). Wnt-induced GSK3 and Zyxin vesicles were quantified in Figure 2E and F. Overexpression of transfected CA-LRP6-GFP lacking the extracellular domain, elicits a potent Wnt signal mediated by the formation of MVBs (Taelman et al., 2010), and caused colocalizarion with endogenous Zyxin (*Figure 2-figure Supplemental S1*). These results indicate that Wnt signaling causes the endocytosis of focal adhesion proteins into the same vesicles that sequester GSK3, suggesting a connection between the Wnt pathway and the focal adhesion signaling pathway.

**Figure 2.**
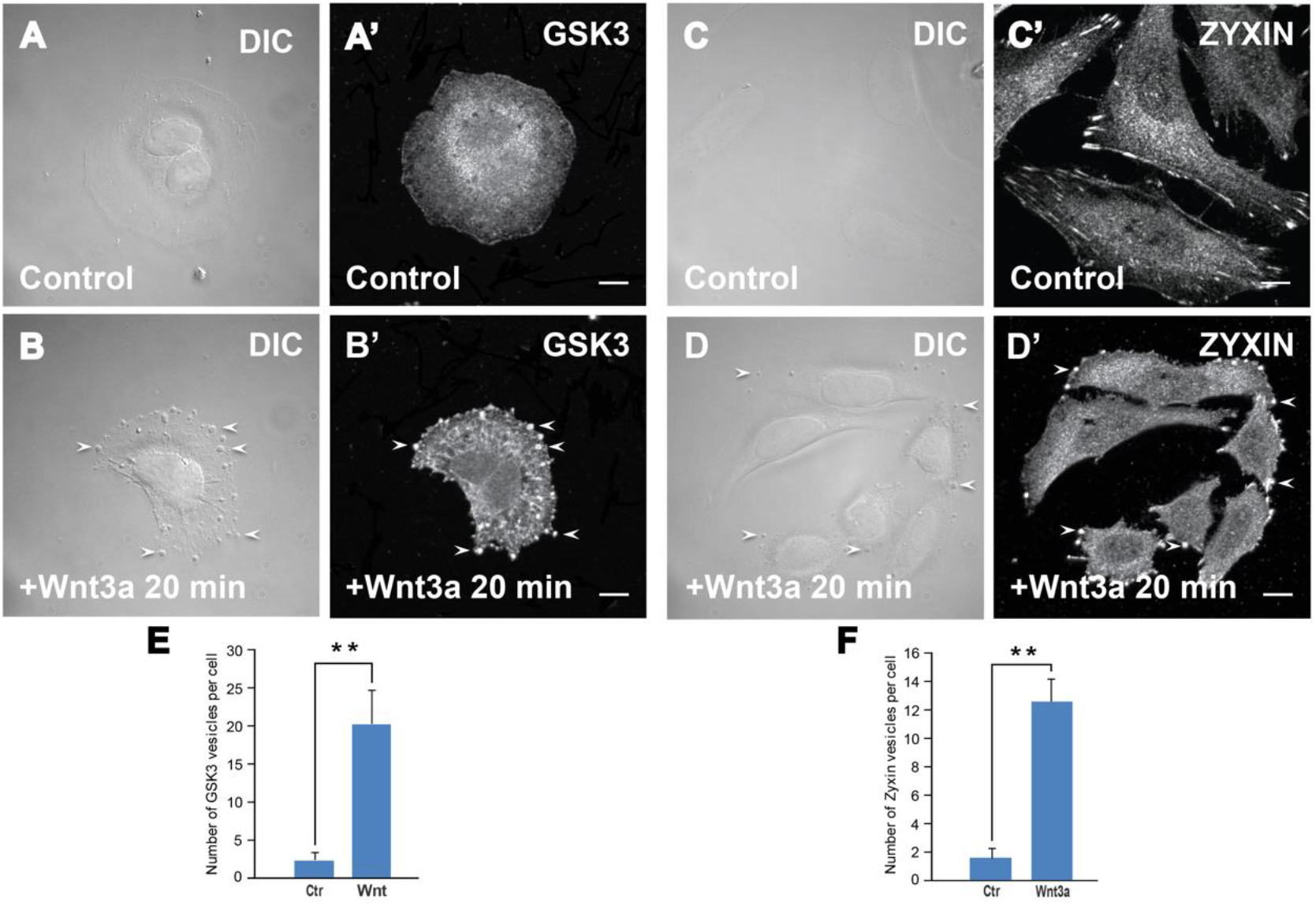
Wnt3a treatment caused the formation of vesicles that sequestered GSK3 and the focal adhesion protein Zyxin. Immunostainings of endogenous GSK3 and Zyxin. (***A****-**B**’*) Fluorescence microscopy images in HeLa cells showing that Wnt treatment (100 ng/ml, 20 min) caused the translocation of GSK3 from the cytosol into vesicles. Note that Wnt3a protein caused the formation of prominent vesicles visible by light microscopy (arrowheads). (***C****-**D**’*) The focal adhesion protein Zyxin is sequestered in vesicles, as GSK3 is, by 20 min Wnt stimulation. Images were generated using a Zeiss Imager Z.1 microscope with Apotome using high magnification. (***E****-**H***) Quantification of the immunofluorescence found in vesicles (see methods). Scale bars, 10 μm. Error bars denote SEM (n ≥ 3) (**p < 0.01). **Figure supplement 1.** Wnt signaling induced by CA-LRP6-GFP relocalizes the Focal Adhesion Zyxin into MVB vesicles.

### Plasma membrane Integrin β 1 is rapidly endocytosed after Wnt treatment

Integrins are constantly endocytosed and recycled back to the plasma membrane through multiple pathways (Moreno-Layseca et al., 2019; Li et al., 2020). Tight regulation of integrin turnover from the cell surface is pivotal to a number of biological processes. This includes cell migration, cytokinesis, and cancer cell invasion and metastasis. Integrins are predominantly endocytosed via clathrin-mediated endocytosis (CME), but other mechanisms such as caveolin and macropinocytosis have been implicated (Gu et al., 2011; Moreno-Layseca et al., 2019).

To examine whether Wnt3a affected endocytosis of ITGβ1, a surface biotinylation assay was performed (Figure 3). HeLa cells were treated with Wnt3a protein on ice or incubated at 37°C for 15 or 30 min before placing on ice (low temperature prevents endocytosis), and the plasma membrane was labeled with non-cell permeable sulfo-NHS-SS-Biotin for 30 min. Biotinylated proteins were pulled-down with Streptavidin-agarose beads followed by a western blot with ITGβ1 antibody. Wnt3a treatment resulted in a rapid depletion of ITGβ1 from the cell surface (Figure 3, compare lane 6 to lanes 7 and 8). Transferrin receptor (TfR), which is recycled independently of the Wnt pathway, was used as a plasma membrane control, and remained unchanged. We conclude that Wnt3a treatment induces acute endocytosis of ITGβ1 from the cell surface.

**Figure 3.**
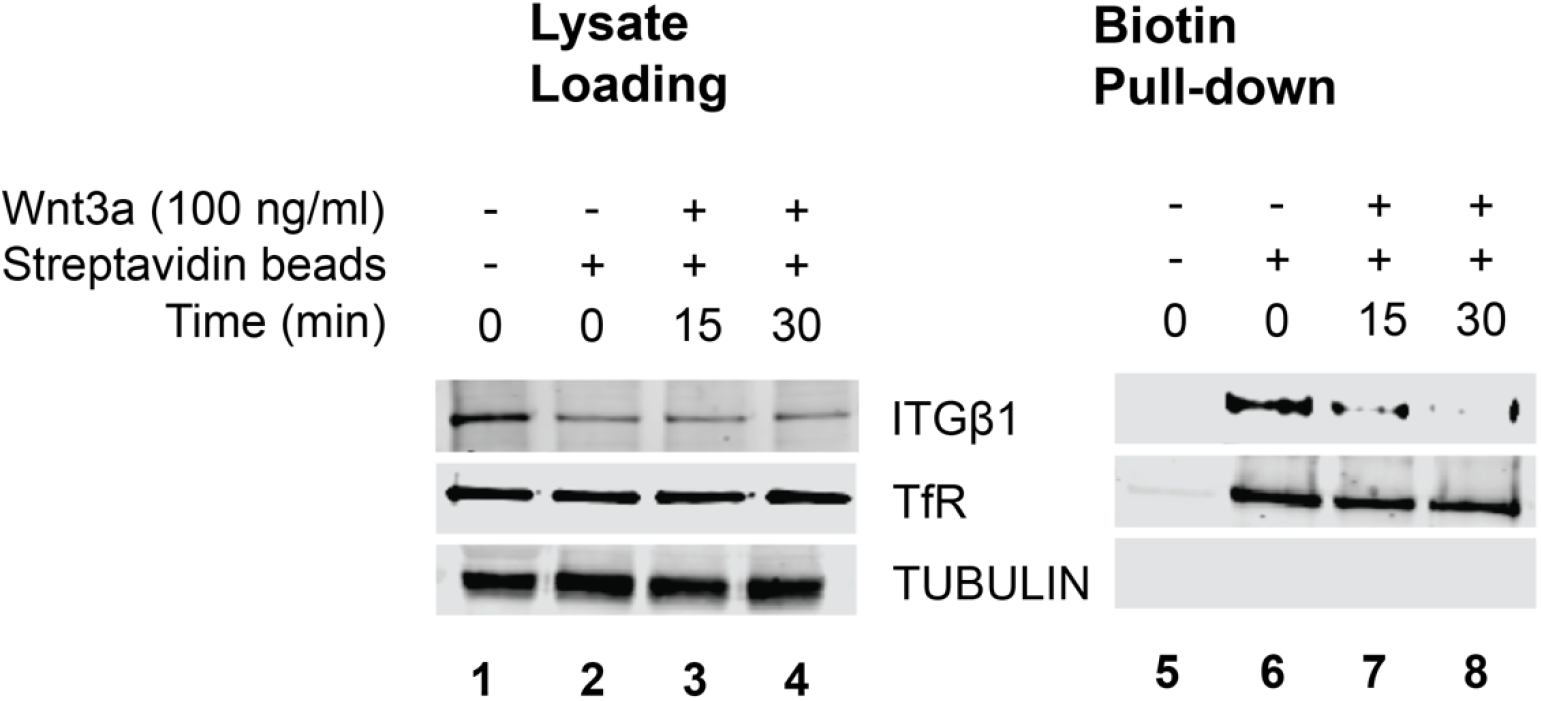
Integrin Beta-1 is rapidly endocytosed by Wnt. Time course (0-30 minutes) of Wnt3a treatment was made in HeLa cells at a permissive temperature for endocytosis. After that, the plasma membrane was labeled with sulfo-NHS-SS-Biotin on ice for 30 minutes. Pull down assay with streptavidin-agarose beads showed that Wnt treatment induced the endocytosis of ITGβ1 after 15 minutes. Lanes 1-4 indicate Hela Cell lysate loading controls, and lanes 5-8 samples after pull down with biotin agarose beads. Cell extracts were analyzed by western blot with ITGβ1 antibody. Note that cell surface ITGβ1 is endocytosed after 15 or 30 min of Wnt3a treatment (lanes 5-8). Transferrin Receptor (TfR) was used as a specificity control that is recycled independently of the Wnt pathway. Tubulin antibodies served as cytoplasmic contamination control. Similar results were obtained in three independent experiments. **Figure supplement 1.** Transcription of genes in the Integrin pathway, such as Talin 1 (TLN1) and ITGβ1 (arrowhead), is reduced in HCC cells mutated in the GSK3 binding site of Axin1.

To investigate whether the sustained activation of the Wnt pathway that occurs in cancer cells also alters focal adhesions, we used the Alexander1 HCC±Axin1 system, which carries a mutation in the GSK3 binding site of Axin1 (Albrecht et al., 2020). RNA-seq genome set enrichment analyses (GSEA) showed that mutation of Axin1 strongly affected the BioCarta Integrin pathway set of 34 genes, leading to a reduction in the transcript levels of Talin 1 (TLN1) and Integrin-β1 when compared to cells in which Axin1 had been reconstituted (*Figure 3-figure supplement 1A*, arrowhead). This result was confirmed by immunostaining of HCC±Axin1 cells on fibronectin-coated slides, which showed that while abundant ITGβ1 focal adhesions were present in cells reconstituted with wild-type Axin1, they were greatly reduced in Axin1 mutant cells (*Figure 4-figure supplement 1B and C*). We conclude that in a cancer cell model, sustained activation of the Wnt pathway by mutation of a tumor suppressor strongly reduces ITGβ1 and focal adhesions at a transcriptional level, in addition to the rapid Wnt-induced clearing of Integrin β-1 by endocytosis.

**Figure 4.**
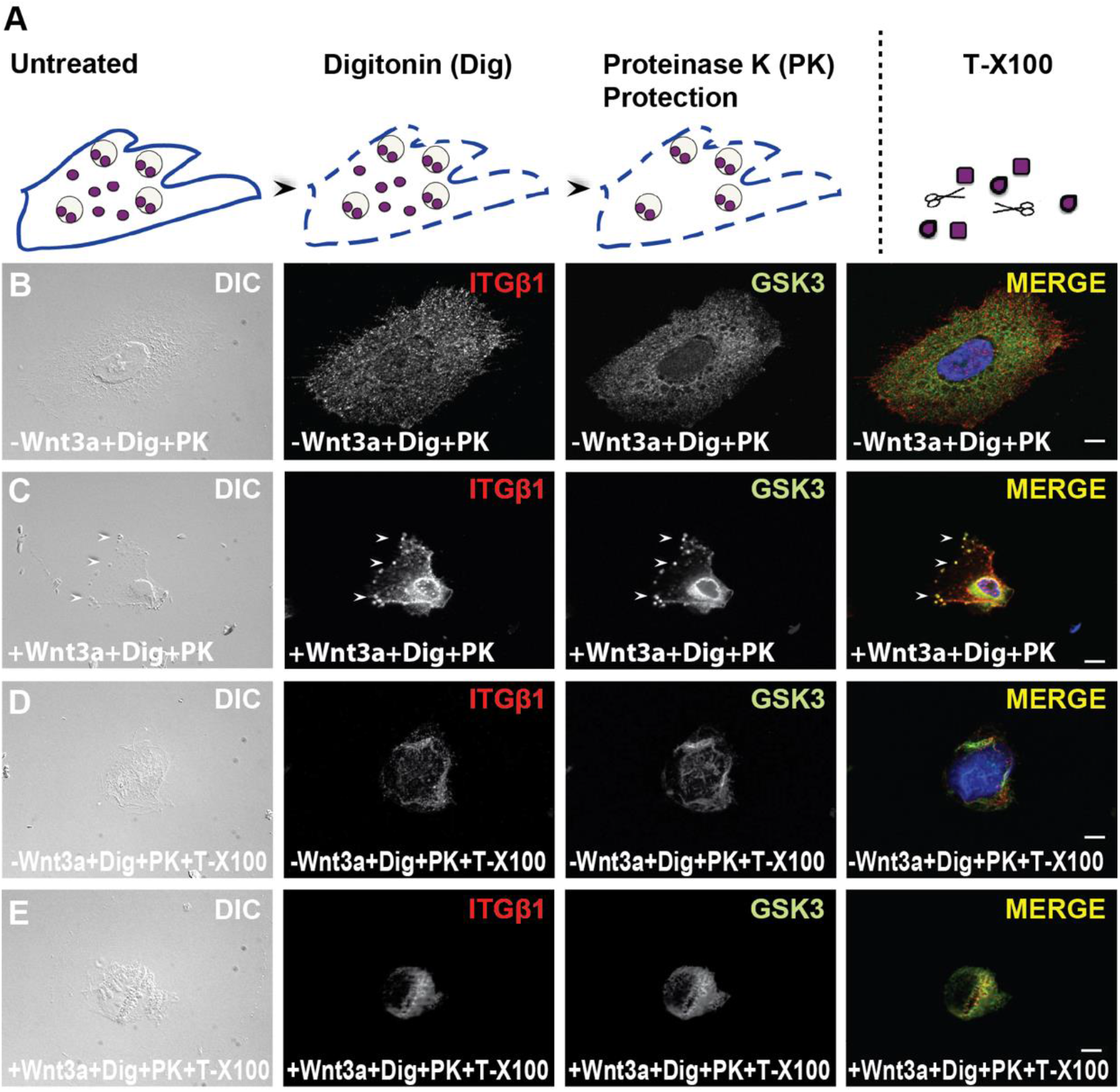
GSK3 and Integrin β-1 are protected from Proteinase K digestion inside membrane-bounded organelles in digitonin-permeabilized cells, but not in the presence of Triton X-100 which solubilizes intracellular membranes. (***A***) Diagram of steps involved in the *in situ* protease protection assay. (***B*** *and **C***) HeLa cells plated on glass coverslips were permeabilized with digitonin, treated with Proteinase K to digest cytosolic proteins, stained with ITGβ1 and GSK3 antibodies, and analyzed by fluorescence microscopy. ITGβ1 and GSK3 were protease-protected within the same vesicles after treatment with Wnt3a protein for 20 min. (***D****-**E***) HeLa cells were treated as described above, except for the addition of Triton, used as control, which dissolves inner membranes and leads to the digestion by exogenous protease of ITGβ1 and GSK3; DAPI labels nuclei. All of the assays were performed in triplicate. Scale bars, 10 μm. **Figure supplement 4.** Phosphorylated Src tyrosine kinase co-localizes into vesicular puncta together with GSK3 20 minutes after Wnt3a addition.

### GSK3 and ITGβ1 are Protected from Protease Digestion inside MVBs after Wnt3a Treatment

The gold standard for determining the localization of a protein inside a membrane-bounded compartment is the protease protection assay (Vanlandingham and Ceresa, 2009). To test whether focal adhesion proteins were translocated inside membrane vesicles after Wnt3a treatment, we used HeLa cells permeabilized with digitonin. Digitonin solubilizes patches of cholesterol-rich plasma membrane but leaves intracellular membranes unaffected (see diagram in Figure 4A). Cells were treated ± Wnt3a, placed on ice, and permeabilized in situ with digitonin (Albrecht et al., 2018). The addition of Proteinase K degraded cytosolic proteins (Figure 4B), but in Wnt-treated cells ITGβ1 and GSK3 were protected from protease digestion inside common vesicles (compare Figure 4C, arrowheads). These puncta corresponded to membrane-bounded organelles, since treatment with 0.01% Triton X-100, which dissolves all intracellular membranes, eliminated the protease protection of ITGβ1 and GSK3 (Figure 4D and E).

We next tested whether another component of the focal adhesion pathway, c-Src, was also endocytosed in ITGβ1-containing vesicles. We chose c-Src because it is an important proto-oncogene Tyrosine kinase, known to regulate the cytoskeleton and the disassembly of focal adhesions in its activated form (Kaplan et al., 1994; Frame et al., 2002). When the proteinase K protection assay was performed using an anti-phospho-c-Src antibody that marks phosphorylated Tyr416 in the activation loop, it was found that Wnt3a treatment also relocalized c-Src into Wnt-induced membrane that contained GSK3.

To determine whether the protease-protected organelles corresponded to MVBs, we used vacuolar protein sorting 4 (Vps4) labeled with GFP. Vps4 is an ATPase component of the ESCRT machinery required in the final stages of pinching off intraluminal vesicles (Gruenberg and Stenmark, 2004). A very useful reagent is provided by a point mutation in the ATP binding site, called VPS4-EQ, which generates a potent dominant-negative protein that prevents MVB formation and Vps4 localization to MVBs (Tejeda-Muñoz et al., 2019). Untreated HeLa cells transiently transfected with wild-type Vps4-GFP showed MVBs that did not colocalize strongly with ITGβ1 focal adhesions (Figure 5A). However, in cells treated with Wnt3a for 20 minutes Vps4-GFP relocated to ITGβ1-containing puncta (Figure 5B, arrowheads). Transfected dominant-negative Vps4-EQ showed that ITGβ1 did not overlap with Vps4-EQ-GFP, both in the absence or presence of Wnt3a, providing a specificity control for MVB localization (Figure 5C and D).

**Figure 5.**
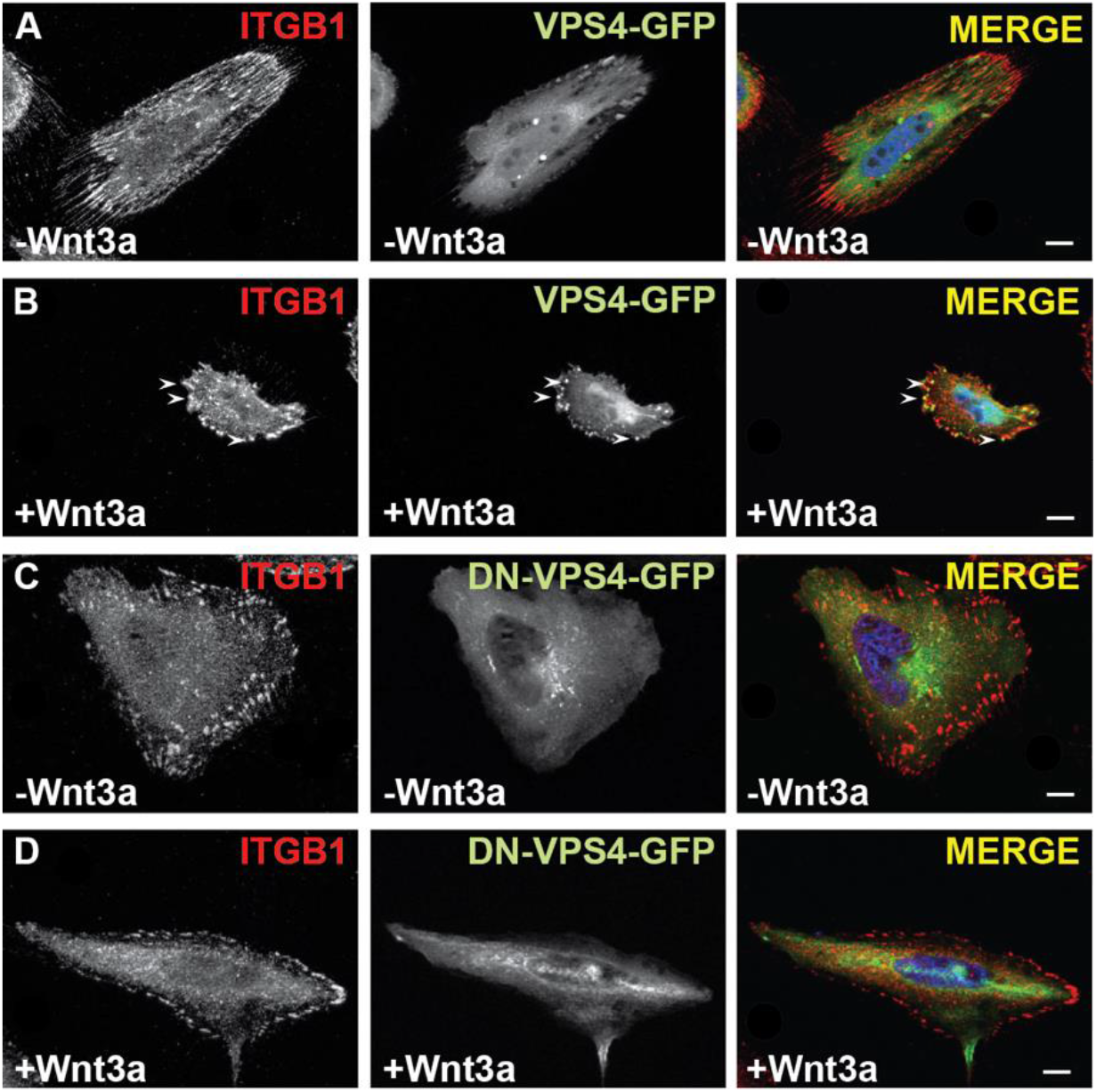
Integrin β-1 co-localizes together with the multivesicular endosome marker Vps4, but not with its dominant-negative point mutant Vps4-EQ. (***A****-**B***) HeLa cells transiently transfected with Vps4-GFP were analyzed using fluorescence microscopy with anti- ITGβ1 antibody. Wnt3a treatment of 20 min caused the re-distribution of ITGβ1 into puncta, known to correspond to MVBs (Taelman et al., 2010), indicated by arrowheads. (***C****-**D***) Overexpression of the Dominant-Negative-Vps4-GFP construct containing a single mutation (EQ) in the ATP binding site blocked the induction of ITGβ1 vesicles by Wnt protein. All assays were performed in triplicate. Scale bars, 10 μm.

We conclude that after short Wnt treatments Integrin β-1 is relocalized from focal adhesions to membrane-bounded organelles that contain GSK3. In addition, the proto-oncogene c-Src, which promotes focal adhesion disassembly, was also translocated into these vesicles. The Wnt-induced puncta corresponded to multivesicular bodies containing the ESCRT marker Vps4.

### ITGβ1 MO reduces Wnt Signaling

To investigate whether ITGβ1 has a cross-talk with Wnt signaling, we used *Xenopus* embryos, which are widely used as a Wnt assay system (Niehrs, 2021). We developed a sensitized system in which microinjection of 0.5 pg of *xWnt8* mRNA into the animal pole of each 4-cell stage blastomere consistently resulted in the complete dorsalization of the embryo, leading to development into radial head structures (Figure 6A and B). We designed an ITGβ1 antisense morpholino (MO) oligonucleotide. When injected on its own, ITGβ1 MO was without phenotypic effect (Figure 6C), but when co-injected together with xWnt8 it reduced dorsalization and allowed for the formation of partial axial structures (Figure 6D). This decrease in Wnt-induced signaling was specific for ITGβ1 depletion, as it was restored by microinjection of human ITGβ1 mRNA that differs in the MO-targeted region (Figure 6E and F). In cultured cells, depletion of ITGβ1 by siRNA (*Figure 6-figure supplement 1A*) or by a DN-ITGβ1 construct (*Figure 6-figure supplement 1B*) (Retta et al., 1998), also reduced Wnt signaling. Taken together, the results indicate that proper cell adhesions via ITGβ1 facilitate signaling by the Wnt pathway.

**Figure 6.**
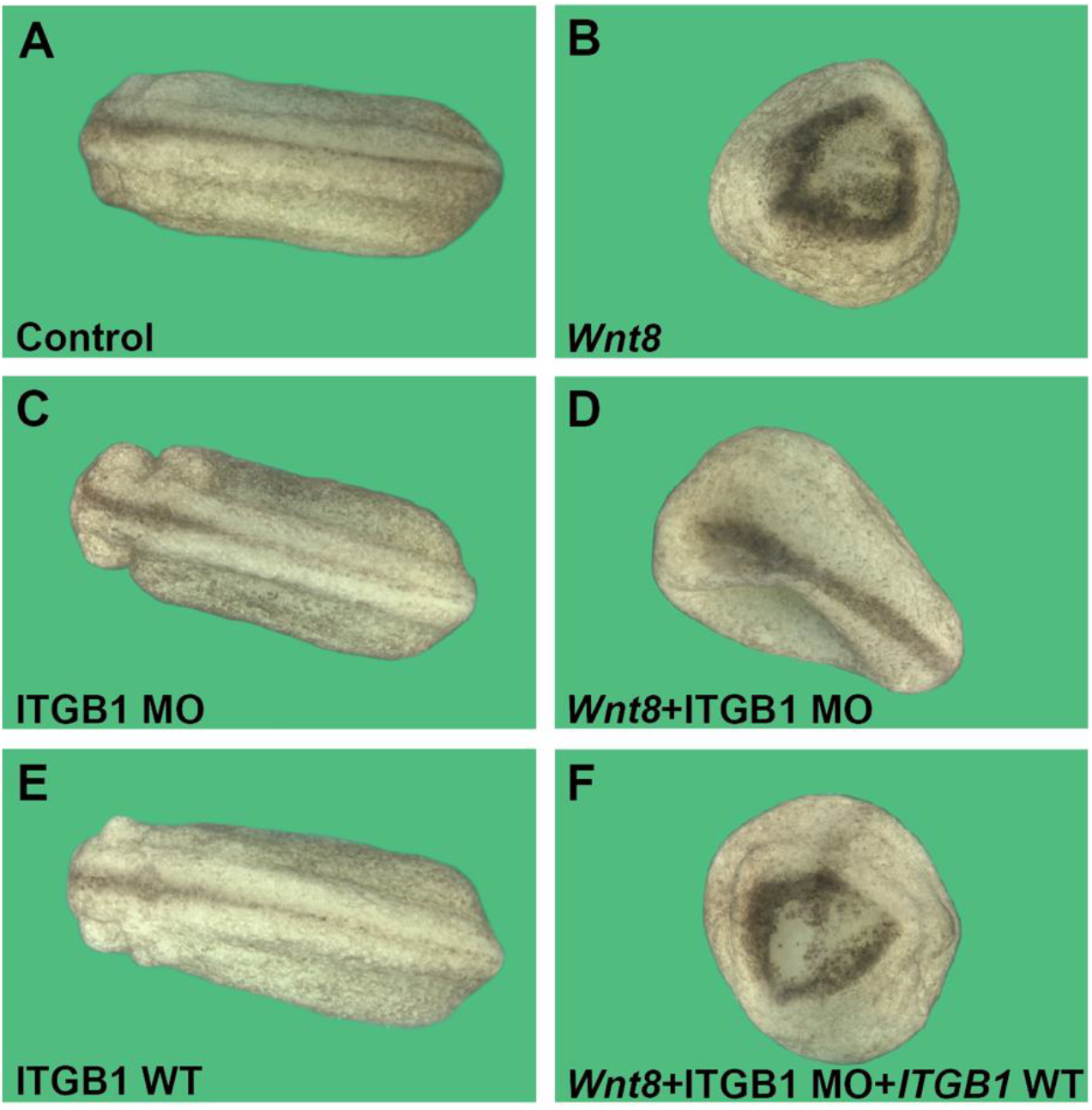
ITGβ1 MO Inhibits Wnt Signaling in a sensitized *Xenopus* embryo assay in which injection of *xWnt8* four times into the animal pole induces a radial dorsalized phenotype. (***A***) control *Xenopus* embryos at early tailbud. (***B***) Embryos injected four times in the animal region at the 4-cell stage with 0.5 pg *Wnt8* mRNA consistently induced a radial head phenotype lacking any trunk development. (***C***) No phenotype was observed with ITGβ1 MO alone. (***D***) Co-injection of xITGβ1 antisense MO consistently inhibited the dorsalizing effects of xWnt8, allowing the formation of partial axial structures. (***E***) microinjection of human *ITGβ1* mRNA was without phenotypic effect. (***F***) the effect of ITGβ1 MO was specific because it was rescued by human *ITGβ1* mRNA not targeted by the MO sequence. The images were taken with an Axio Zoom V.16 Stereo Zoom Zeiss at low magnification. Similar results were obtained in five independent experiments. Numbers of embryos analyzed were as follows A=140; B=128; C=132; D=125; E=129; F=124 (5 independent experiments). Scale bar, 500 μm. Figure 6 **supplement 1.** Canonical Wnt signaling is inhibited by ITGβ1 depletion in mammalian cells.

## Discussion

The main finding in this study is that a crosstalk exists between canonical Wnt signaling and focal adhesions. As shown in Figure 7, binding of Wnt to its receptors Lrp6 and Frizzled causes a local inhibition of GSK3 activity, an enzyme that normally inhibits macropinocytosis mediated by the cellular actin machinery in the leading edge lamellipodium (Tejeda-Muñoz et al., 2019; Albrecht et al., 2020). The receptor complex is rapidly endocytosed into late endosomes/MVBs that fuse with lysosomes which become more acidic degrading macropinocytosed macromolecules (Albrecht et al., 2021). The sequestration of GSK3 inside the endosomal compartment via the ESCRT machinery is required for sustained canonical Wnt signaling (Taelman et al., 2010; Vinyoles et al., 2014; Tejeda-Muñoz et al., 2019). Our attention was drawn to focal adhesions by a study showing that a Wnt pathway component, Dishevelled, associated with membrane vesicles that localized to the end of actin cytoskeletal cables (Capelluto et al., 2002), which is also the location of focal adhesions. Primary fibroblasts rearrange the cytoskeleton within 20 min of Wnt3a addition, and the focal adhesion marker Vinculin became internalized (Figure 1). In HeLa cells, Wnt addition caused the formation of prominent vesicles visible by DIC brightfield; these vesicles colocalized with GSK3 and the focal adhesion marker Zyxin (Figure 2). Integrins are transmembrane proteins that anchor the actin cytoskeleton to the extracellular matrix. Consisting of α and β subunits, there are 24 possible heterodimers, yet the majority share a common β-1 subunit, leading us to focus on ITGβ1 (Moreno-Layseca et al., 2019). Cell surface biotinylation studies showed that ITGβ1 protein was endocytosed within 15-30 min of Wnt3a treatment (Figure 3, lanes 6-8). In situ protease protection studies showed that after Wnt signaling ITGβ1 became relocalized to membrane-bounded organelles that were the same ones that sequestered GSK3 (Figure 7). These vesicles were specifically labeled by the MVB marker Vps4 and therefore corresponded to MVBs. An important regulator of focal adhesion disassembly, the proto-oncogene c-Src (Moreno-Layseca et al., 2019), was also relocalized inside Wnt-induced membrane-bounded organelles (*Figure 4-figure supplement 1*). These experiments suggest that, unexpectedly, Wnt-induced endocytosis results in the translocation of multiple focal adhesion components into late endosomes.

**Figure 7.**
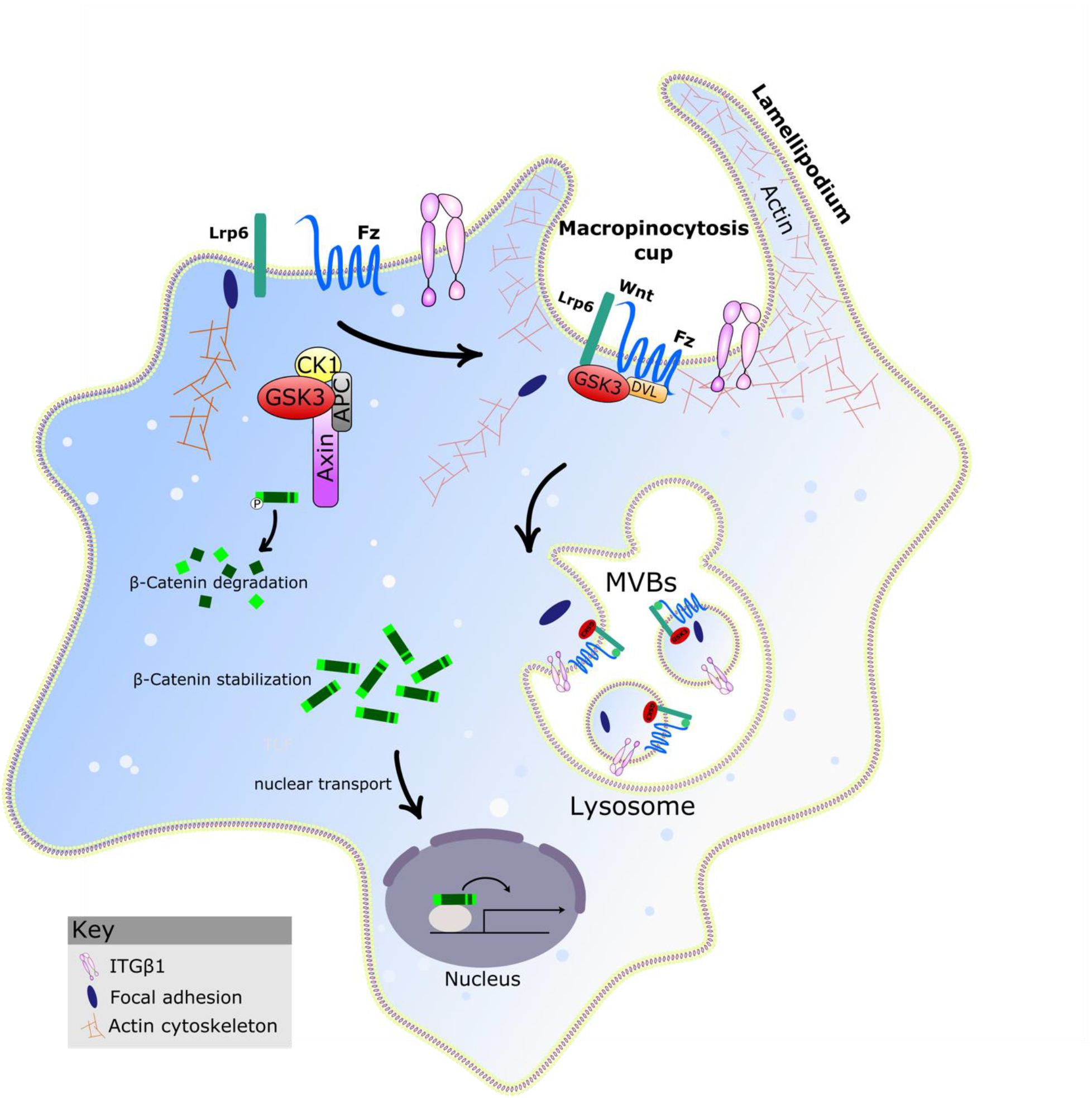
Model of the Endocytosis of the Focal Adhesion and ITGβ1 by Wnt. After Wnt treatment, Lrp6/Fz/Wnt/GSK3 signalosomes, the master regulator of cell adhesion ITGβ1 and other focal adhesion components are endocytosed by macropinocytic cups in an actin-driven process. Macropinocytosis is required for canonical Wnt signaling. Sequestration of GSK3 in MVBs is necessary for the stabilization of β-catenin that mediates the transcriptional activity of canonical Wnt.

Previous indications existed in the literature suggesting interactions between focal adhesion and Wnt signaling, which are two major pathways of cellular transduction. For example, it has been reported that extracellular matrix stiffness causes increased transcription of Wnt1 through the activation of the integrin-Focal Adhesion Kinase (FAK) pathway (Du et al., 2016). The focal adhesion protein Kindlin-2, was found to directly bind to β-catenin and TCF4 forming a transcriptional complex that enhances Wnt signaling in tumors (Yu et al., 2012). Recent studies on cell polarization and migration have shown that key elements in the Wnt pathways such as GSK3 are essential to cell polarity. GSK3 influences cell migration as one of the regulators of the spatiotemporally controlled dynamics of the actin cytoskeleton, microtubules, and cell-to-matrix adhesions (Sun et al., 2009). In this regard, it has been reported that GSK3 phosphorylates FAK, reducing its activity and consequently impeding cell spreading and migration (Bianchi et al., 2005). In contrast, other experiments indicate that GSK3 promotes cell spreading by phosphorylating paxillin, a focal adhesion constituent (Cai et al., 2006). It is known that the Tyrosine kinase c-Src, which participates in focal adhesion disassembly, also binds to and phosphorylates Dishevelled 2, dampening Wnt transcriptional signals (Yokoyama and Malbon, 2009).

The ESCRT machinery has been shown to regulate the transit of c-Src from endosomes to the plasma membrane, promoting the turnover of cell-ECM adhesions during cell migration (Tu et al., 2010). In our experiments, the ESCRT machinery was shown to be involved in the endocytosis of ITGβ1 as well as that of c-Src. In an interesting recent study, it was found that macropinocytosis serves as a novel mechanism that shuttles actin to the leading edge for lamellipodium expansion during neural crest migration (Li et al., 2020).

Although there are multiple examples of interactions between Wnt and focal adhesions in the literature, the results presented here differ in that we discovered that focal adhesions are rapidly translocated into MVB vesicles after Wnt signaling. Furthermore, using a sensitized *Xenopus* embryo assay in which microinjection of *xWnt8* mRNA caused completely radial embryos consisting of heads and no tails, we found that depletion of ITGβ1 attenuated the effects of Wnt overexpression (Figure 6). This suggests that in the context of the embryo, proper cell adhesion through focal adhesions facilitates cell-cell Wnt signaling. Since Wnt downregulates focal adhesions, and Integrin depletion dampens Wnt signaling, the two pathways might establish a negative feedback loop that limits the duration of Wnt signals.

The transition from single-celled to multicellular organisms marked the emergence of complex life forms. Understanding molecular and cellular mechanisms of key cell signaling pathways in both development and disease has been a major focus of research. The Wnt and Integrin signaling pathways are very ancient and were already present in sponges (Brower et al., 1997; Loh et al., 2016). Wnt signaling drives a cell growth program that can become inappropriately activated in cancer. Wnt and cell adhesion are often active in the same developmental processes and crosstalk between them should result in reciprocal regulation. Knowing how Wnt signaling and focal adhesions cooperate will improve our understanding of embryonic development and tumorigenesis. Cancer therapies in the past have focused on blocking the uncontrolled cell division of primary tumors. However, almost all cancer deaths are due to metastatic cancer, not primary tumors. We uncovered here a new regulation model by which Wnt signaling changes cell adhesion through endocytosis of focal adhesion components. It has been proposed that focal adhesion restoration strategies could be useful in the treatment of metastatic cancer (Griffith et al., 2005; Li et al., 2016; Chaturvedi et al., 2014; Ramirez et al., 2011). Understanding how macropinocytosis, Wnt, and focal adhesions intersect could lead to new targets in cancer treatment.

## Materials and Methods

### Antibodies and Reagents

Antibodies against the focal adhesion protein Zyxin (ab50391, 1:200), Vinculin (ab129002, 1:200), Phalloidin (ab176759, 1:1000), and the serine threonine kinase GSK3 (ab93926, 1:4000) were obtained from Abcam. ITGβ1 antibody was obtained from Proteintech (12594-1-AP, 1:100), while anti-γ-Tubulin antibody was obtained from Sigma (T6557, 1:3000). Secondary antibodies for immunostaining (1:500) were from Jackson ImmunoResearch Laboratories, Inc (715166150, 711545152). Secondary antibodies coupled to Infrared Dyes (IRDye 680 (926– 68072) and IRDye 800 (926–32213) at 1:3000 (LI-COR) were used for western blots and analyzed with the LI-COR Odyssey system. Primers for cloning human DN-ITGβ1 into pCS2 c- terFlag were Inte-Forward: GGCGGATCCACCATGAATTTACAACCAATTTTCTG, Inte- Reverse: GGCATCGAT TATCATTAAAAGCTTCCATATCAG. Primers for cloning human ITGβ1-WT into pCS2 Inte-WT- Reverse: GGCATCGATTTTTCCCTCATACTTCGGA. Pools of 3 target-specific ITGβ1 (sc-35674) and control siRNAs (sc-37007) were obtained from Santa Cruz Biotechnology. *Xenopus laevis* ITGβ1 antisense MO, sequence 5’GTGAATACTGGATAACGGGCCATCT3’, was designed with the help of GeneTools.

### Tissue Culture and Transfection

HeLa (ATCC, CRL-2648), HEK-293BR (BAR/Renilla), NIH 3T3, Alexander cells (RRID:CVCL_0485), and HCSF cells were cultured at 37 ° C in 5% CO2 atmosphere in DMEM (Dulbecco’s Modified Eagle Medium), supplemented with 10% fetal bovine serum, 1% glutamine, and penicillin/streptomycin. The cells were seeded to maintain a cell density between 20%-60% and experiments were performed when cells reached a confluence of 70-80%. Cells were transfected by Lipofectamine 3000. DNA constructs were added to cells and incubated overnight, with a fresh change of media after 12-16 hours. Cells were transferred to 2% of fetal bovine serum 6-12 hours after transfection before treatment with Wnt3a (Preprotech Cat# 315-20) protein at 100 ng/ml.

### Immunostainings

HeLa or HCSF cells were plated on glass coverslips and then directly transfected, or alternatively transfected with a later split onto coverslips. Coverslips were acid-washed and treated with fibronectin (10 μg/ml for 30 min at 37°C, Sigma F4759) to enhance cell spreading and adhesion. Cells were fixed with 4% paraformaldehyde (Sigma #P6148) 15 min, permeabilized with 0.2% Triton X-100 in phosphate-buffered saline (PBS; Gibco) for 10 min, blocked with 5% BSA in PBS for 1 hour. Primary antibodies were added overnight at 4°C. The samples were washed three times with PBS, and secondary antibodies were applied for one hour at room temperature. After three additional washes with PBS, the coverslips were mounted with Fluoroshield Mounting Medium with DAPI (ab104139). Immunofluorescence was analyzed and photographed using a Zeiss Imager Z.1 microscope with Apotome.

### Western Blots

Cell lysates were prepared using RIPA buffer (0.1% NP40, 20 mM Tris/HCl pH 7.5), 10% Glycerol, together with protease (Roche #04693132001) and phosphatase inhibitors (Calbiochem #524629).

### Cell Surface Biotin Labeling

HeLa cells were incubated with Wnt3a for 0, 15, and 30 minutes at 37 °C. The cells were placed ice and washed three times with ice-cold PBS (Fisher Scientific), and then labeled with non-cell permeable sulfo-NHS-SS-Biotin (1 mg/ml) for 30 min using rotary agitation at 4°C (Ding et al., 2018). The cells were washed three times with ice-cold quenching solution (50 mM Glycine in PBS, pH 7.4), and with ice-cold PBS. Cell lysates were prepared using a RIPA buffer and incubated with Streptavidin-agarose beads (Thermo Scientific) using end-over-end agitation at 4°C overnight. An aliquot of the original cell lysate was saved for input control. The resin was washed 3 x with TNE buffer (Sigma) and then an equal volume of 2 x SDS loading sample buffer was added, and heated at 95°C for 5 min. The samples were electrophoresed by polyacrylamide gel electrophoresis (PAGE), and transferred to a nitrocellulose membrane using a semi-dry transfer apparatus (1 hour at 15 V). The membranes were blocked (TBS with skimmed milk 5%) for an hour. Later the membrane was incubated with primary (overnight) and secondary antibodies (60 minutes), and blots were developed using the LiCor Odyssey system.

### Protease Protection Assay

Cells were plated on glass coverslips and after 24 hours were treated with or without Wnt3a for 20 minutes. They were placed on ice, permeabilized with digitonin (6.5 μg/mL) for 30 minutes, and incubated with Proteinase K (1 μg/mL) for 10 minutes (Albrecht et al, 2018). As a control, the addition of Triton X-100 (0.01%) was used to dissolve all cell membranes. Samples were then analyzed by immunofluorescence.

### *Xenopus* Embryo Microinjection and In Situ Hybridization

*Xenopus laevis* embryos were fertilized *in vitro* using excised testis. Staging was as described (Nieuwkoop and Faber, 1967). *In vitro* synthesized mRNAs were introduced into embryos by microinjection using an IM 300 Microinjector (Narishige International USA, Inc) 4 times. pCS2- hITGβ1, was linearized with NotI and transcribed with SP6 RNA polymerase using the Ambion mMessage mMachine kit. Embryos were injected in 1x MMR and cultured in 0.1x MMR (Albrecht et al, 2020). To study the role of ITGβ1 in the Wnt pathway, we designed an ITGβ1 antisense MO against *Xenopus* ITGβ1. Injection of 4 nl of 0.3 mM MO four times into the animal pole of 4-cell stage embryos was without phenotypic effect. However, ITGβ1 MO interfered with the dorsalization caused by the radial injection of *Wnt8* mRNA (0.5 pg). This effect was specific to ITGβ1 as axial development could be rescued by co-injection of 100 pg of human *ITGβ1* mRNA.

### Animal Cap Cell Culture

Animal caps were dissected at early blastula stage 8.5 to 9 in 1 x MMR solution, washed 3 times, and cells cultured in L-15 medium containing 10% Fetal Calf Serum diluted to 50% with H20 (Smith and Tata 1991) on fibronectin-coated coverslips for 12-18 h. When mounting with coverslips, a drop of Anti-fade Fluorescence Mounting Medium-Aqueous, Fluoroshield was added and microscopic examination of RFP and GFP was performed. Image acquisition was performed using a Carl Zeiss Axio Observer Z1 Inverted Microscope with Apotome.

### Library preparation and Illumina sequencing

RNA was isolated with the RNAeasy Plus kit (QIAGEN) from 500-900 x 10^3^ cells. Four independent cultures of matched HCC±Axin1 cells were used. The quality and concentration of the extracted material was assessed using the RNA assay on a 2200 TapeStation (Agilent Technologies): all the samples had a RIN score higher than 8. Libraries were constructed with the KAPA mRNA HyperPrep kit (Roche Sequencing, cat# KK8580/ 08098115702) with unique dual-indexed adapters according to the manufacturer’s protocol. Final library QC qPCR quantitation was performed using the D1000 assay on a 2200 TapeStation (Agilent Technologies). After concentration normalization, libraries were pooled and sequenced on a NovaSeq 6000 (Illumina) SP lane (2x150).

### Sequencing data processing

After sequencing, demultiplexed reads (Illumina bcl2fastq) were processed to remove the adapters using fastp (Chen et al., 2018). Trimmed reads were aligned against the GRCh38 genome using STAR (v2.7.4.a). The counts assigned to each gene were combined into a matrix and subject to differential expression analysis using DESeq2. The complete RNA-seq data reported in this paper have been deposited in GEO, Accession Number GSE193381

### Heatmap

Normalized raw counts (DESeq2) were used as input for unbiased heatmap visualization using the R package heatmap with the following options: scale = ’row’, cutree_rows = 4, cluster_cols = TRUE, cluster_rows = TRUE. Only genes defined as members of the Integrin pathway (BioCarta) are shown.

### Statistical Analyses and Image Quantification

The data are expressed as means and standard errors of the mean (SEM). Statistical analysis of the data was performed using the Student t-test. A P value of <0.01 was considered statistically significant. The number of vesicles in the immunofluorescence was quantified from the DIC light microscope channel using the ImageJ software and a computer-assisted particle analysis tool. Individual channels were given thresholds with MaxEntropy. The individual vesicles were separated using the “binary watershed” function and vesicles were counted using the “analyze particles” function, with particle size 0.2–5 μm and circularity –1. More than 25 cells were counted per experiment. They were numbered, outlined, and then copied to a spreadsheet in order to perform the statistical analysis.

## Acknowledgments

We are grateful to D. Geissert for technical assistance, H. Coller for human corneal fibroblasts, R. Moon for BAR reporters, J. Monka and Y. Ding for comments on the manuscript, and M. F. Domowicz for help with the illustrations.

## Author contributions

N.T.M designed research and wrote the paper; N.T.M., Y.M., P.S., M.M., M.P., and E.M.D.R. performed research and analyzed data.

## Supplementary Materials

**Figure 1 supplement 1.**
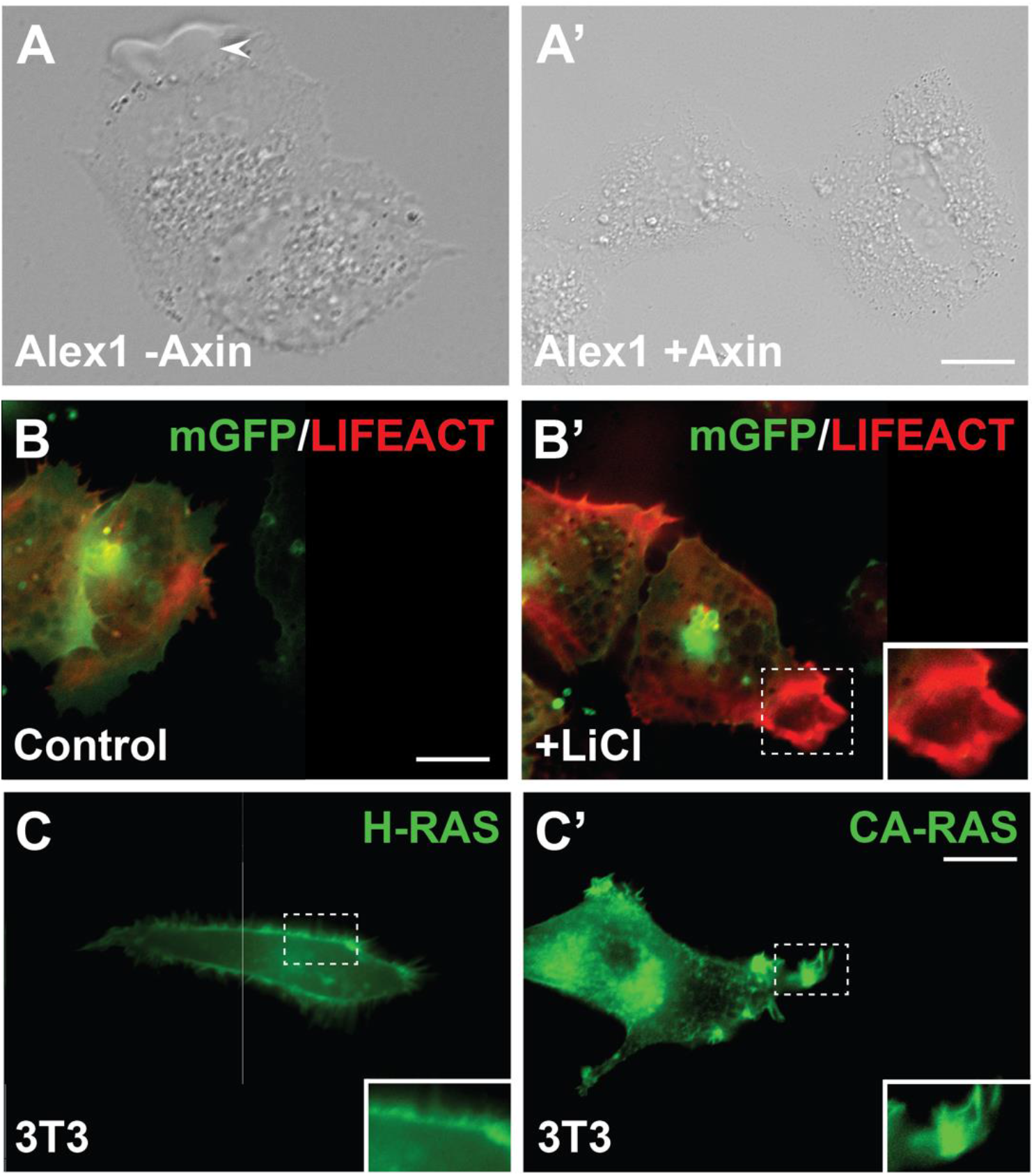
Macropinocytosis in different cellular models; see also Videos 1, 2 and 3. (***A*-*A*’**) Alexander1 hepatocellular carcinoma cells showing that mutation of Axin1 causes an increase in ruffling in the plasma membrane cell membrane (arrowhead) indicative of macropinocytosis activity. (***B***-***B***’’) LiCl treatment triggers the formation of large macropinocytosis cups (stippled square) in animal cap cells injected with Lifeact (in red) and membrane-GFP (in green) mRNAs in *Xenopus* animal cap cells plated on fibronectin. (***C*-*C*’**) Activated HRas-GFP (G12V mutation), but not wild-type HRas-GFP, induced sustained macropinocytosis (stippled square), used here as a control for macropinocytic activity, in transfected 3T3 cells. Shown are still images of Videos 1, 2, and 3.

**Figure 2 supplement 1.**
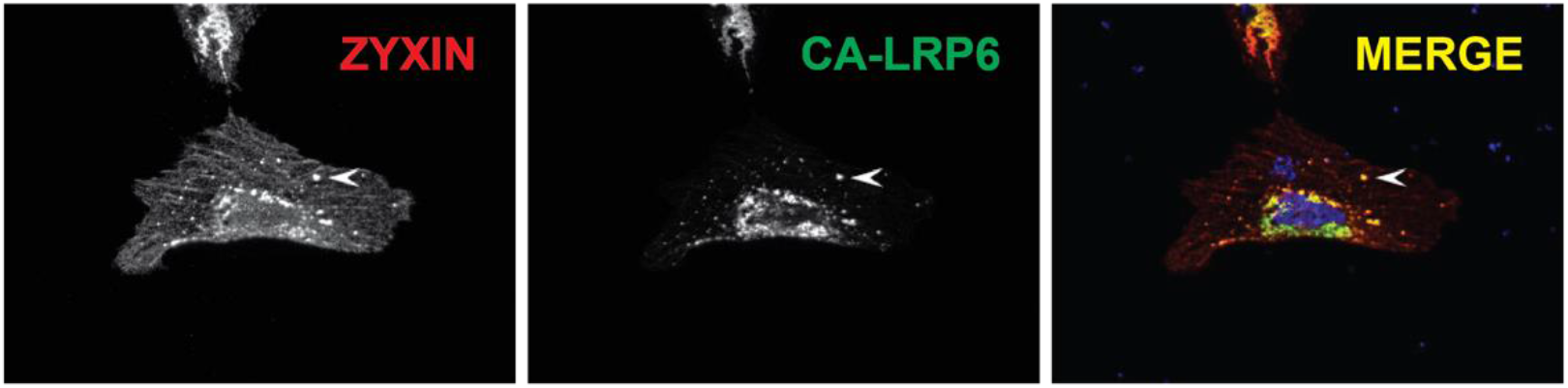
Wnt signaling induced by CA-LRP6-GFP relocalized the Focal Adhesion Zyxin into MVB vesicles. Fluorescence microscopy images of HeLa Cells transiently transfected with CA-LRP6-GFP and immunostained for Zyxin. Note that Zyxin co-localized with CA-LRP6-GFP vesicles (arrowhead), which are known to correspond to multivesicular endosomes (Taelman et al., 2010). All assays were performed in triplicate.

**Figure 3 supplement 1.**
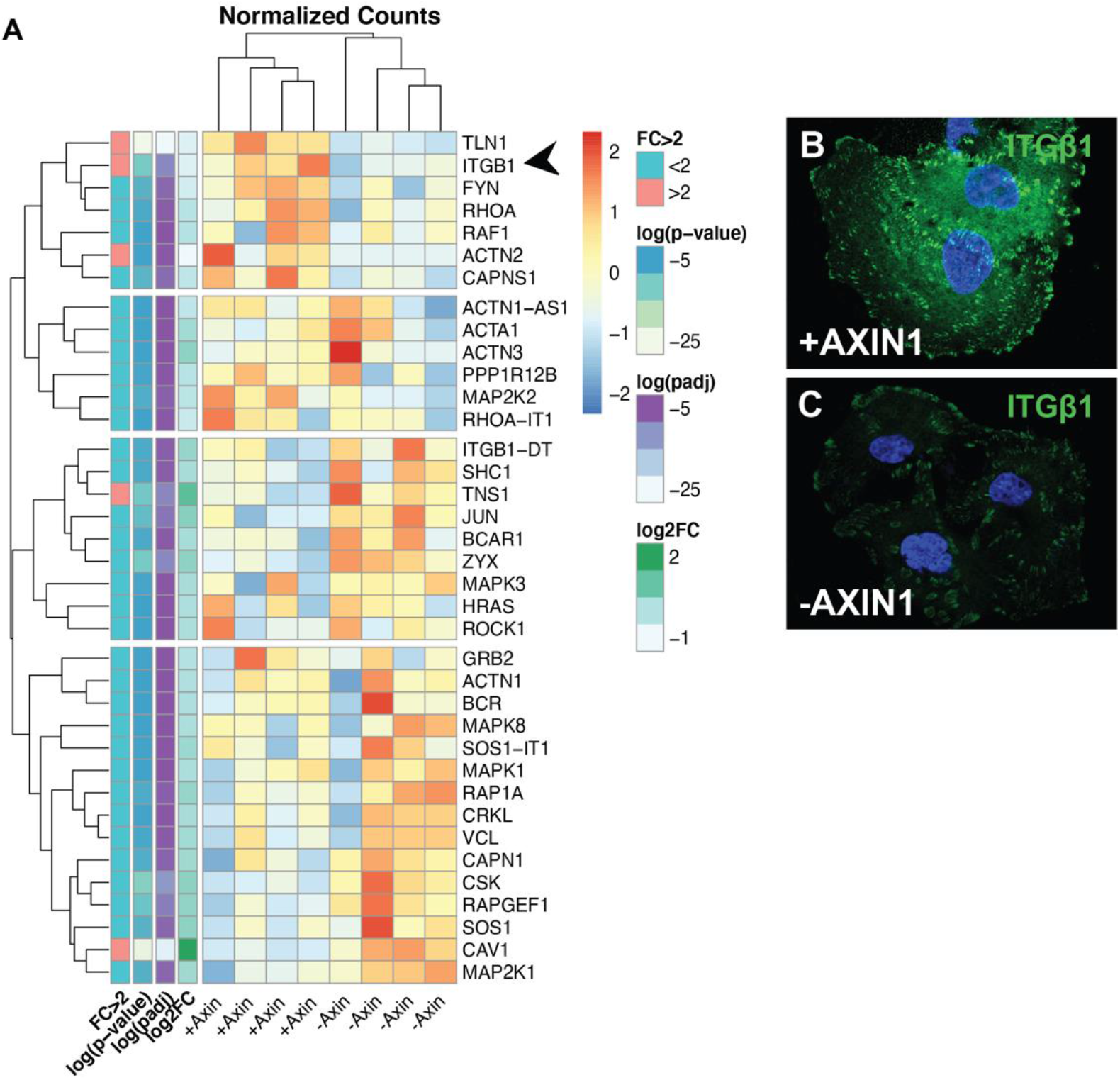
Transcription of genes in the Integrin pathway, such as Talin 1 (TLN1) and ITGβ1 (arrowhead), is reduced in HCC cells mutated in the GSK3 binding site of Axin1. We developed an HCC model using Alexander1 cells (ATCC CRL-8024) mutated in Axin1. Using retroviral VSVG transfection and hygromycin B clonal selection, we restored Flag-tagged Axin1 (Addgene #24928) expression in these cells (Albrecht et al., 2020). (***A***) Heatmap from RNAseq data of Alexander HCC cells ± Axin1. Note that depletion of the tumor suppressor Axin dramatically affects expression of the BioCarta Integrin pathway (and triggers macropinocytosis in these cells, *Video 1*). (***B*, *C***) Axin1 mutation resulted in a striking decrease in ITGβ1- containing focal adhesions.

**Figure 4 supplement 1.**
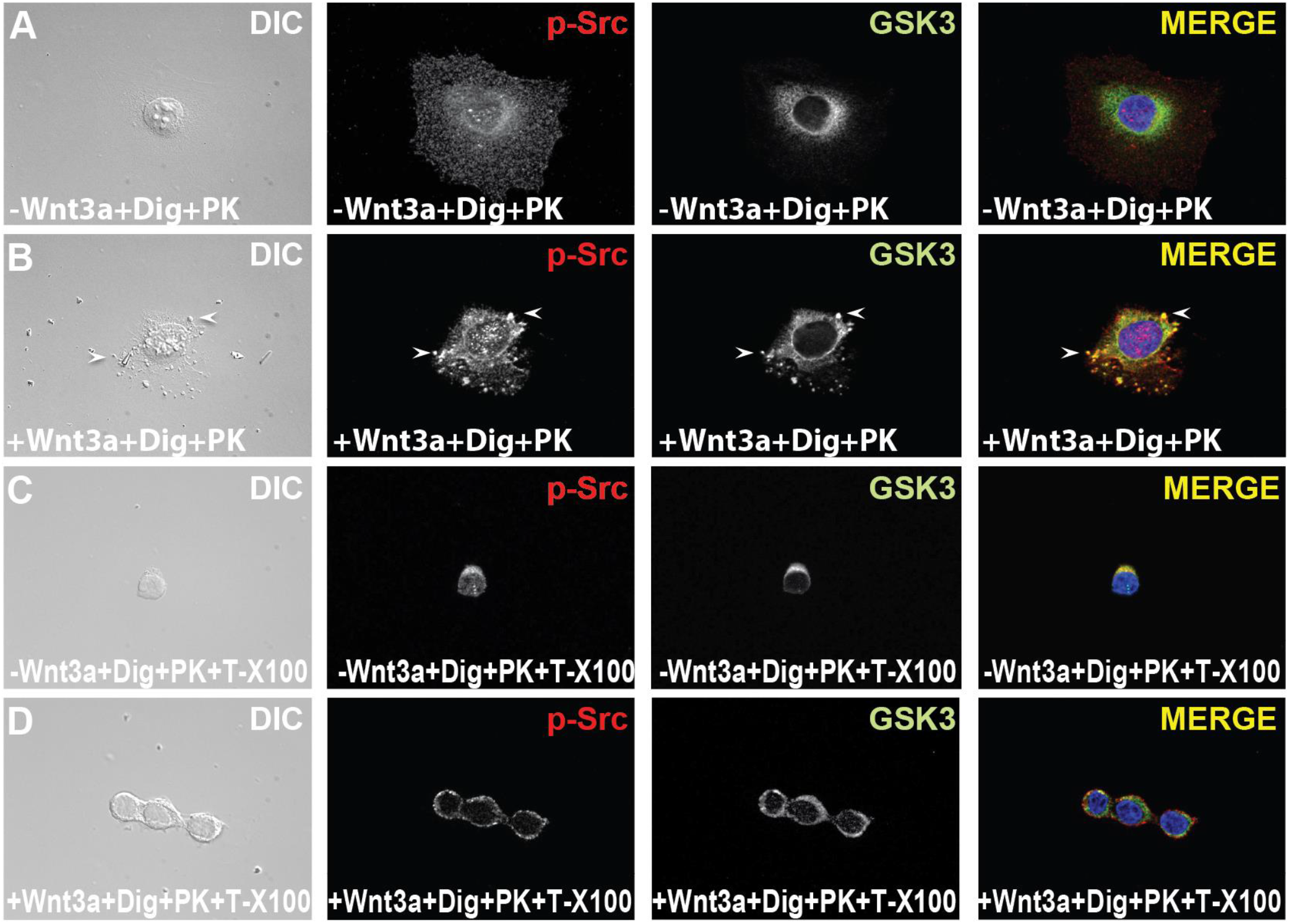
Phosphorylated c-Src tyrosine kinase translocates into vesicular puncta together with GSK3 after 20 minutes of Wnt3a addition. *In situ* protease protection experiment showing that p-Src and GSK3-containing vesicles were protected from proteinase K digestion in digitonin-permeabilized cells, but not in the presence of 0.01% Triton X-100, which solubilizes all membranes. All assays were performed in triplicate.

**Figure 6 supplement 1.**
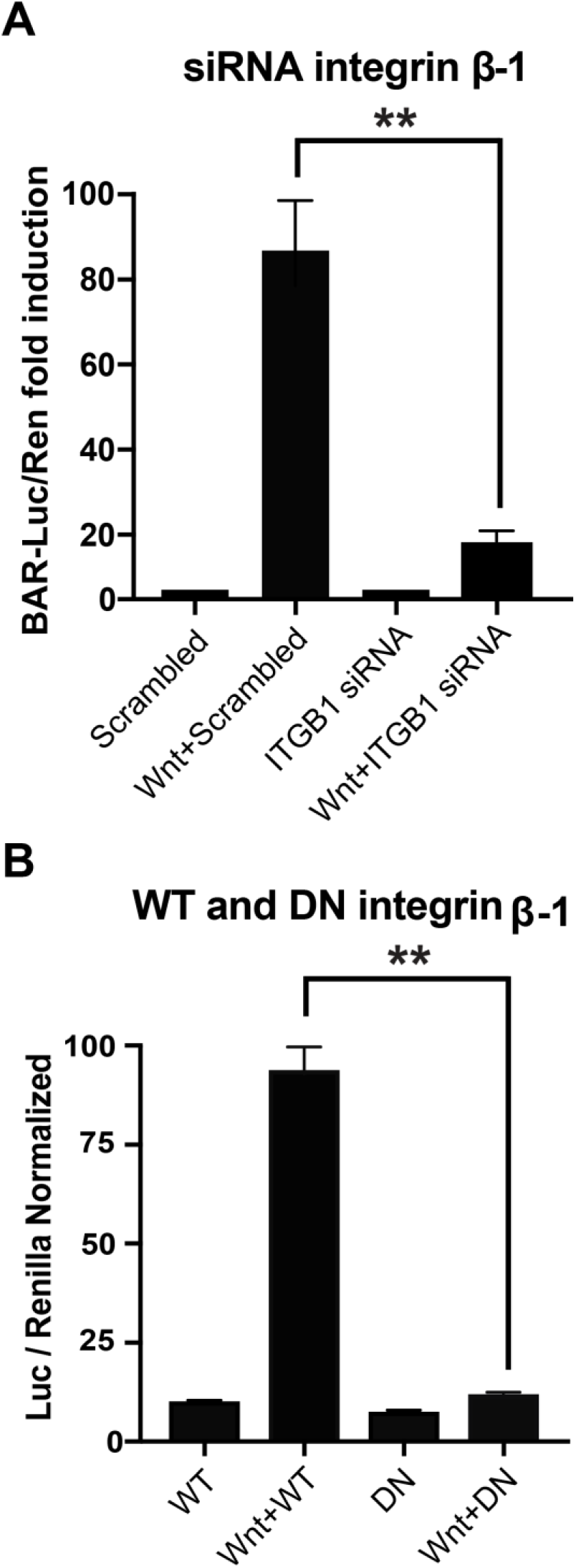
Canonical Wnt signaling is inhibited by ITGβ1 depletion in mammalian cells. (***A***) HEK-293 cells were co-transfected with β-catenin reporter genes and either with scrambled siRNA or with ITGβ1 siRNA. After 72 hours post-transfection, cells were incubated with Wnt3a ligand (100 ng/ml) for 12 hours. Note that the transcriptional activity of β-catenin was inhibited by ITGβ1 siRNA, but not by scrambled siRNA. (***B***) Tranfection of Dominant- Negative Integrin β-1 (DN-ITGβ1), but not wild-type ITGβ1, also resulted in the inhibition of Wnt signaling. Firefly luciferase activity was normalized with respect to the activity of Renilla luciferase in each sample. Error bars denote SEM (n ≥ 3) (**p < 0.01).

## Supporting Information Videos

**Video 1.** Mutation in Axin1 causes extensive membrane ruffling and macropinocytosis in HCC cells. DIC from Alexander1 cells showing a wave-like movements of the lamellipodia and ruffles in the plasma membrane characteristic of macropinocytosis. Axin1 mutation leads to constitutive macropinocytosis. Cells were plated on fibronectin-coated 35 mm Petri dishes with glass bottoms (#0 cover glass, Cell E&G: GBD00003-200) for 12-18 h. The movie was generated with a Zeiss Observer.Z1 microscope equipped with a high magnification Apotome.2. Image acquisition was 40 frames in 20 min. For video editing, the Adobe Premiere Pro CC 2020 software was used.

**Video 2.** Mimicking Wnt with the GSK3 inhibitor LiCl triggers macropinocytic cup formation in cultured *Xenopus* animal cap cells. F-actin tracer LifeAct (Gonagen) (in red) and mGFP mRNAs (in green) were injected in frog embryos 4 times into the animal pole at the 4 cell-stage. Animal caps were dissected at early blastula stage 9 in 1 x MMR solution, washed 3 times, and cells plated in L-15 medium containing 10% Fetal Calf Serum diluted to 50% with H20 onfibronectin-coated 35 mm Petri dishes with glass bottoms for 12-18 h. Control cells were filmed for 20 min. Then LiCl (40 mM final) was added and filmed for an extra 20 min. Note that the leading edge of the cell changes with LiCl treatment, forming macropinocytosis cups. Fluorescence filters were controlled by Axiovision 4.8 software and saved in this program. Video was generated with the same methods as described in Video 1.

**Video 3.** Activated Ras-GFP triggers macropinocytosis. mEGFP HRas (Addgene# 18662) and mEGFP HRas G12V (Addgene# 18666) were transfected in 3T3 cells and filmed the next day for 20 min. Note that in the controls the cell membrane lacks the plasma membrane ruffling typical of macropinocytosis. In cancer cells, activated Ras is known to induce sustained macropinocytosis. Movies were generated with the same conditions as described in Video 1.

